# Biophysical basis of phage liquid crystalline droplet-mediated antibiotic tolerance in pathogenic bacteria

**DOI:** 10.1101/2022.12.13.520211

**Authors:** Jan Böhning, Miles Graham, Suzanne C. Letham, Luke K. Davis, Ulrike Schulze, Phillip J. Stansfeld, Robin A. Corey, Philip Pearce, Abul K. Tarafder, Tanmay A. M. Bharat

**Affiliations:** Structural Studies Division, MRC Laboratory of Molecular Biology, Francis Crick Avenue, Cambridge CB2 0QH, United Kingdom; Sir William Dunn School of Pathology, University of Oxford, Oxford OX1 3RE, United Kingdom; Department of Mathematics, University College London, London WC1H 0AY, United Kingdom; Institute for the Physics of Living Systems, University College London, London WC1E 6BT, United Kingdom; Cell Biology Division, MRC Laboratory of Molecular Biology, Francis Crick Avenue, Cambridge CB2 0QH, United Kingdom; School of Life Sciences & Department of Chemistry, University of Warwick, Coventry, United Kingdom; Department of Biochemistry, University of Oxford, Oxford OX1 3QU, United Kingdom

## Abstract

Inoviruses are abundant filamentous phages infecting numerous prokaryotic phyla, where they can symbiotically promote host fitness and increase bacterial virulence. Due to their unique properties, inoviruses have also been utilised in biotechnology for phage display and as models for studying phase behaviour of colloidal rods. Inoviral phages secreted by bacteria can self-assemble into liquid crystalline droplets that protect bacterial cells in biofilms from antibiotics, however, factors governing the formation of such droplets and the mechanism of antibiotic protection are poorly understood. Here, we investigate the structural, biophysical, and protective properties of liquid crystalline droplets formed by *Pseudomonas aeruginosa* and *Escherichia coli* inoviral phages. We report a cryo-EM structure of the capsid from the highly studied *E. coli* fd phage, revealing distinct biochemical properties of fd compared to Pf4 phage from *P. aeruginosa*. We show that fd and Pf4 form liquid crystalline droplets with diverse morphologies governed by the underlying phage particle geometry and biophysics, rather than their surface biochemical properties. Finally, we show that these morphologically diverse droplets made of either phage can protect rod-shaped bacteria from antibiotic treatment, despite differing modes of association with cells. This study advances our understanding of phage assembly into liquid crystalline droplets, and provides insights into how filamentous molecules protect bacteria from extraneous molecules under crowding conditions, which are found in biofilms or on infected host tissues.

## Introduction

Viruses that infect bacteria and archaea, known as phages, are among the most common biological entities on Earth (Edwards and Rohwer, 2005, Hatfull and Hendrix, 2011). Filamentous bacteriophages, belonging to the family *Inoviridae,* are one of the major categories of phages, being pervasive in prokaryotes across Earth’s biomes (Roux et al., 2019). Inoviruses consist either of a linear single-stranded (Tarafder et al., 2020) or circular single-stranded DNA genome (Xu et al., 2019) bound by a filamentous array of an *α*-helical capsid protein. These rod-shaped inoviruses are typically ∼60-70 Å in diameter, but, depending on their genome size, their length can vary from one to a few microns, as it is proportional to the size of the encapsidated genome. Typically, inoviruses do not lyse cells (Marvin and Hoffmann-Berling, 1963) and either integrate into the host cell genome as prophages (seen for phages Pf4, CTX*ϕ* and MDA*ϕ*), or are expressed by the host cell episomally, i.e, not integrated into the bacterial genome (for fd, M13 and Pf1 phages) (Hatfull and Hendrix, 2011).

The first inovirus identified, named fd, was reported in 1963 as a DNA-containing phage with an unusual filamentous morphology infecting *Escherichia coli* (Marvin et al., 2014, Marvin and Hoffmann-Berling, 1963). Fd has since become a model for filamentous phage structure, infection, and assembly (Marvin, 1998, Marvin et al., 2014), as well as having found use in numerous biotechnological applications, including the first described instance of phage display (Smith, 1985), and as a model for examining colloidal-rod phase transitions (Dogic and Fraden, 2001). Despite its importance in multiple areas of research, past structural studies have reported markedly differing atomic structures of the fd phage capsid (protein pVIII in fd) using fibre diffraction (Marvin et al., 2006, Marvin et al., 1994), solid-state nuclear magnetic resonance (ssNMR, (Zeri et al., 2003, Marvin et al., 2006)) and 8 Å-resolution electron cryomicroscopy (cryo-EM, (Wang et al., 2006)), meaning that the structure of the fd virion remains controversial. Thus far, only two high-resolution cryo-EM structures of filamentous phage capsids have been solved; the first from Ike, which like fd, is a member of the class I type of invoviral bacteriophages with pentameric (C5) capsid symmetry (Xu et al., 2019); and second from Pf4, a class II inoviral bacteriophage with C1 symmetry, which is expressed as a prophage in *P. aeruginosa* biofilms (Tarafder et al., 2020).

A unique property of inoviral phages including fd and Pf4 is that they can self-assemble to form ordered but dynamic liquid crystalline droplets under crowding conditions outside the cell. Within droplets, laterally associated phages are orientationally aligned, but not regularly ordered as in a crystal (Lekkerkerker and Tuinier, 2011). Due to their monodisperse nature, filamentous phages have been used to study phase behaviour of rod-like molecules, with fd having been employed for this purpose for more than half a century (Lapointe and Marvin, 1973, Nakamura and Okano, 1983, Tang and Fraden, 1995, Tomar et al., 2007, Barry et al., 2009, Fiester et al., 2011, Lagerwall et al., 2014). More recently, studies have shown that filamentous phages are highly expressed in bacterial biofilms formed by the human pathogen *P. aeruginosa* (Whiteley et al., 2001). One such phage, called Pf4, is integrated into the *P. aeruginosa* genome, where it is highly upregulated by bacteria upon the switch to a biofilm lifestyle. Pf4 was subsequently found to form liquid crystalline droplets in the biopolymer-rich extracellular matrix of biofilms, where it encapsulates bacterial cells, forming a protective barrier, allowing them to tolerate antibiotic treatment (Secor et al., 2015, Tarafder et al., 2020). The presence of filamentous molecules in a viscous environment is a hallmark of all biofilms (Flemming and Wingender, 2010), however, how these molecules are able to bestow special properties to bacteria including antibiotic protection is unknown.

In this study, to determine the mechanism of protection conferred by phage liquid crystalline droplets, we studied the biochemical and biophysical parameters influencing liquid crystalline droplet formation by inoviral phages. We compared the atomic structures, droplet-forming characteristics, and bacterial encapsulation properties of fd and Pf4 phages on *E. coli* and *P. aeruginosa* cells. Using cryo-EM, we report a 3.2 Å-resolution structure of the intact, native fd capsid, settling a long-standing debate about its structure. Using optical microscopy and electron cryotomography (cryo-ET), we then show that fd and Pf4 form droplets with diverse morphologies, governed by the underlying phage geometry rather than on the biochemical traits of the phages. We show that even though droplets of fd and Pf4 are morphologically diverse and have distinct modes of association with bacterial cells, both can protect rod-shaped bacteria against antibiotic treatment. Our results show how assemblies of filamentous molecules, which are enriched in viscous environments such as bacterial biofilms or sites of infection in hosts, can closely associate with bacterial cells, modulating their response to external stresses, such as antibiotics.

## Results

### Atomic structure of the fd phage capsid from cryo-EM

To study the biochemical properties of the fd phage capsid, we generated phage preparations using previously established procedures (Methods). We observed ∼64 Å wide phage particles on cryo-EM grids (Figure S1A). Using helical reconstruction, a 3.2 Å-resolution map of the fd phage capsid was obtained, which allowed derivation of an atomic model of the capsid (Figures 1A-C and S1B-E, Movie S1 and Table S1). In agreement with previous predictions (Marvin, 1966), the atomic model reveals a pentameric (C5) subunit arrangement of the pVIII capsid protein, which forms a single *α*-helix containing 50 residues. The helix is terminated at the N-terminus of the mature protein by a proline residue (P6), and residues 1-5 of the capsid protein are disordered (Figure 1C). Due to its helical symmetry, capsid protein monomers stack almost vertically to form a highly symmetrical arrangement along the helical axis (Figure 1B). The capsid proteins interact predominantly via a hydrophobic interaction network (Figure 1D), which include the residue Y21 that was shown previously to induce structural instability (Tan et al., 1999) and hence mutated in several studies to methionine (Zeri et al., 2003, Marvin et al., 2006, Blanco et al., 2011). In our specimen of wild-type fd phage, despite retaining Y21, there was no significantly lowered resolution detected at this location in the structure, even though local resolution varied slightly within other parts of the capsid array (Figure S1E).

**Figure 1:**
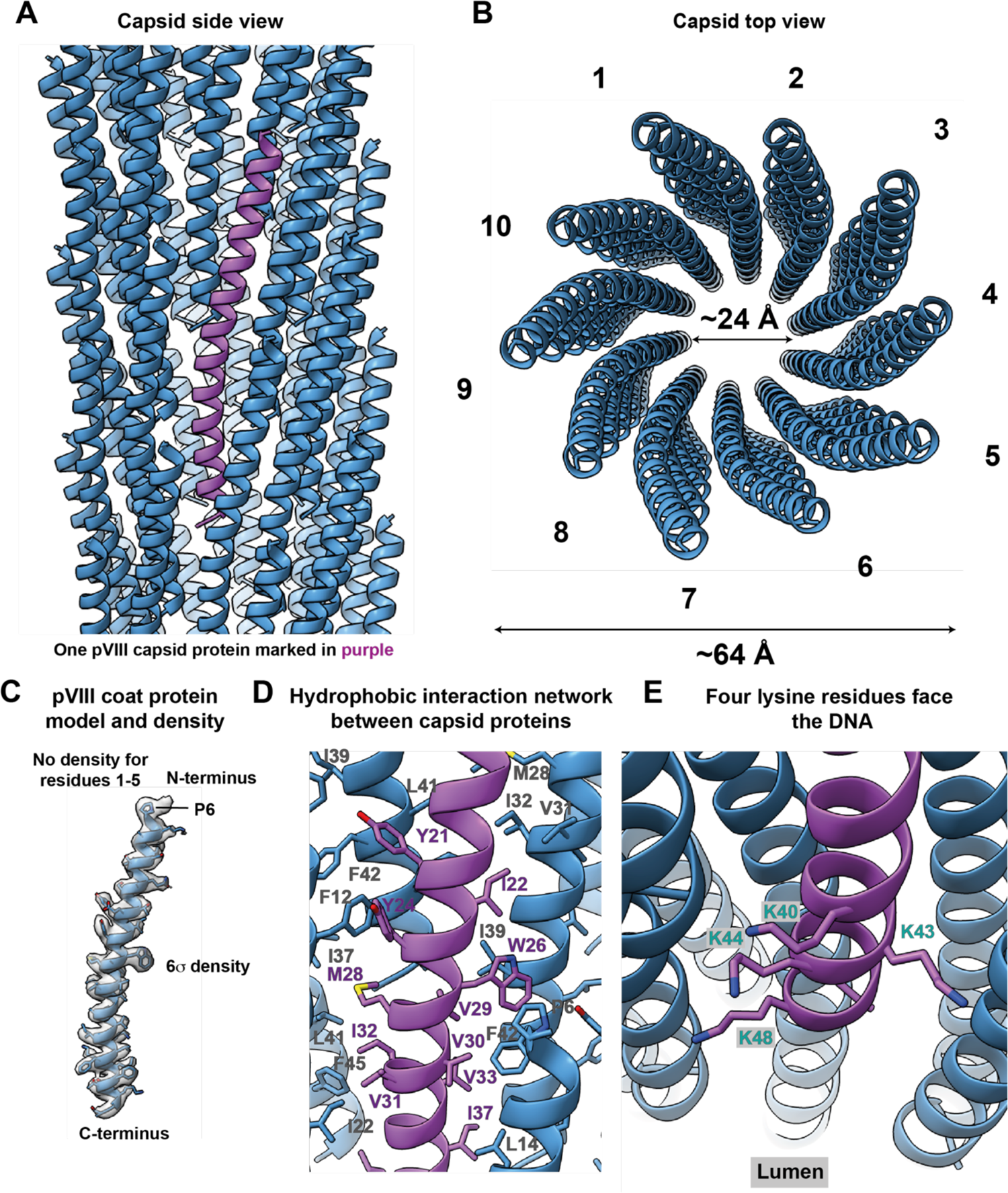
Cryo-EM structure of the fd bacteriophage capsid at 3.2 Å-resolution. **A)** Side view of the fd capsid (ribbon depiction) with a single pVIII subunit highlighted in purple. **B)** Top (perspective) view of the capsid shows a 24 Å-wide lumen. The vertical interactions of capsid proteins results in 10 protofilament-like stacks observed from the top. **C)** Atomic model of pVIII with the cryo-EM density shown as an isosurface at 6 *σ* away from the mean value. **D)** Hydrophobic interaction networks within the capsid, with interacting hydrophobic residues marked. **E)** Four lysine residues near the C-terminus extend into the capsid lumen.

The C-terminus of the capsid protein, which faces the inner lumen of the cylindrical phage, interacting with the genomic DNA, has four positively charged lysine residues exposed to the lumen (Figure 1E). These lysines presumably bind to the negatively-charged DNA phosphates, as also seen in other phages (Xu et al., 2019, Tarafder et al., 2020). Clear density for the DNA was not resolved in our fd cryo-EM map (Figure S1F), in agreement with previous studies on class I inoviral bacteriophages (Figure S2), where an unfeatured central DNA density was reported (Xu et al., 2019). Comparison of our fd capsid structure with previous structural models of fd shows that while the structure of the *α*-helical capsid subunit pVIII has been well-approximated in some structural models, the arrangement of subunits into the overall capsid architecture was imprecise in all (Figure S3).

### Comparison of Class I fd capsid structures with the Class II phage Pf4

Fd is an archetypal class I inoviral bacteriophage that infects *E. coli*, but does not integrate into its genome (Hay and Lithgow, 2019). Structural comparison with Pf4 (Tarafder et al., 2020), a class II inoviral bacteriophage with a prophage lifestyle, shows a slightly larger capsid size (fd: 50 residues, Pf4: 46 residues) and lumen diameter (fd: 24 Å, Pf4: 22 Å, Figure 2). At the N-termini, both phages contain acidic amino acid residues, leading to a negatively charged outer surface (Figure 2B-C). In the case of fd, negatively charged residues are mostly located in the disordered N-terminus (sequence AEGDD), while the negative, outer residues in Pf4 are ordered (Figure 2C). Compared with Pf4, the fd capsid has an increased positive charge facing the inner lumen with four basic residues present at the C-terminus, compared to two in the case of Pf4 (Figure 2D). A higher positive charge-density might be required to sequester a circular single-stranded (ss)DNA of fd, which has been previously suggested (Marvin, 1966), compared to the linear ssDNA of Pf4 (Tarafder et al., 2020), which would require less positive charges from the capsid to stabilise the negative charges of the DNA phosphate backbone.

**Figure 2:**
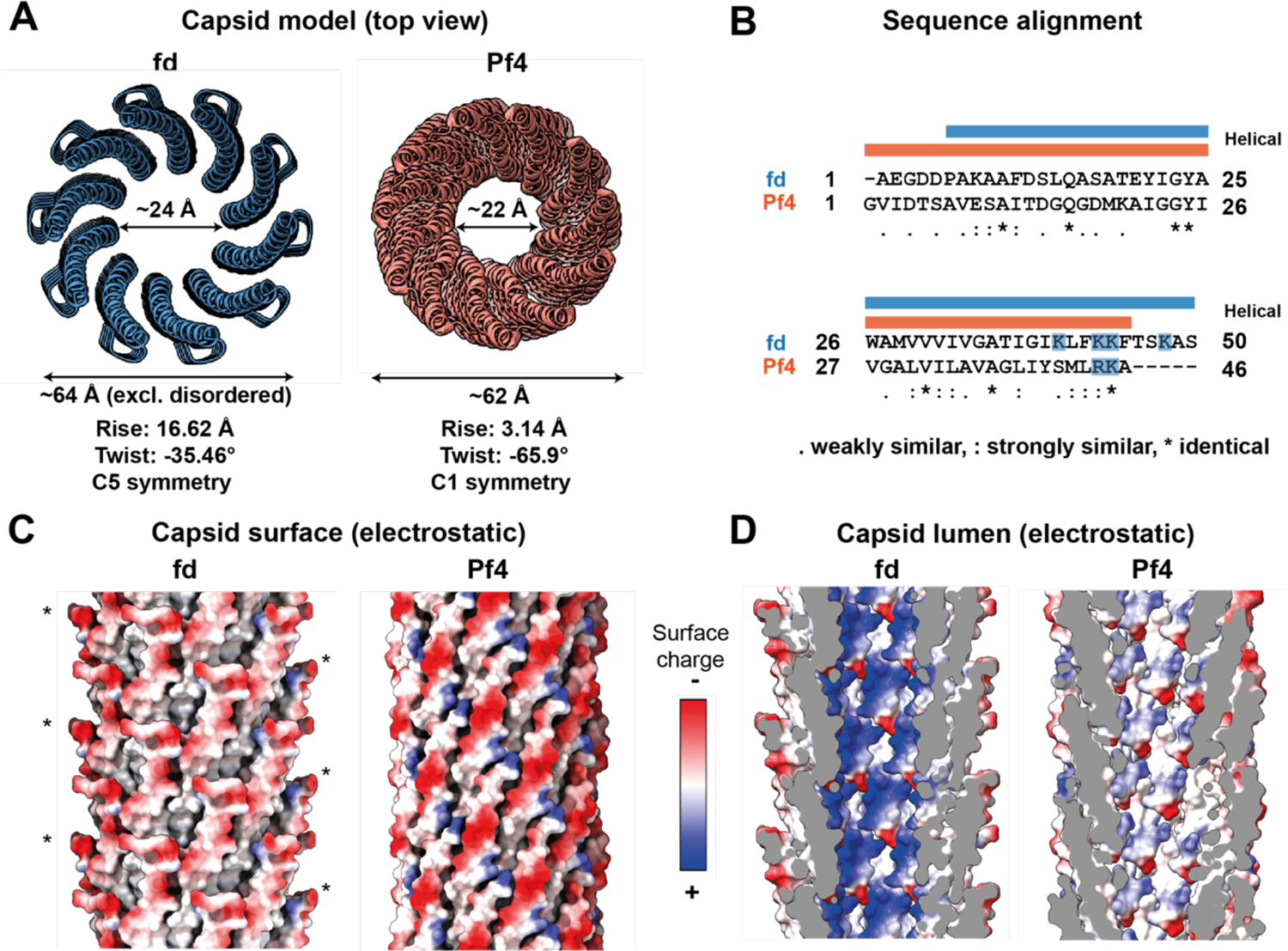
Comparison of the cryo-EM structures of the class I fd phage (this study) and the class II Pf4 phage (PDB 6TUP). **A)** Orthographic top view of fd and Pf4 capsids shown as ribbon diagrams. Flexible residues at the fd N-termini have been modelled to allow comparison with Pf4, where all residues are ordered in the cryo-EM structure. **B)** Clustal Omega sequence alignment of the major capsid protein of fd versus Pf4. Basic residues extending into the lumen are marked in blue. Helical regions are marked above, with fd in blue and Pf4 in salmon. **C)** Electrostatic surface of the capsid. Disordered residues in fd (1-5) were modelled (*) to allow for a more accurate comparison of electrostatic surfaces. **D)** Sliced side view of the capsid lumen depicting electrostatic charge distribution at the luminal surface.

To probe the structure further, we performed atomistic molecular dynamics (MD) simulations of fd and Pf4 phage capsids to examine the characteristics of the lumen and outer residues in a solvated state bound to ions. Comparing simulations of fd and Pf4 capsids, performed in 0.15 M salt (NaCl), where both capsids were stable, we found an increased accumulation of negatively charged chloride ions in the fd lumen (containing four lysine residues) compared with Pf4 (containing only two positively charged residues, Figures 3 and S4). Since our simulations did not contain ssDNA, these increased negatively charged ions in the fd lumen support our expectation from our cryo-EM structures that DNA arrangement in the class I inoviral phage fd (containing circular ssDNA) is different to that in the class II inoviral phage Pf4 (containing linear ssDNA). Also notably, root mean square fluctuation (RMSF) values show flexibility in the N-terminal residues in both phages (Figure S4), with the fd N-terminus significantly extending away from the capsid, confirming the flexible nature of this region, in agreement with the lower local resolution detected in our cryo-EM density (Figure S1E).

**Figure 3:**
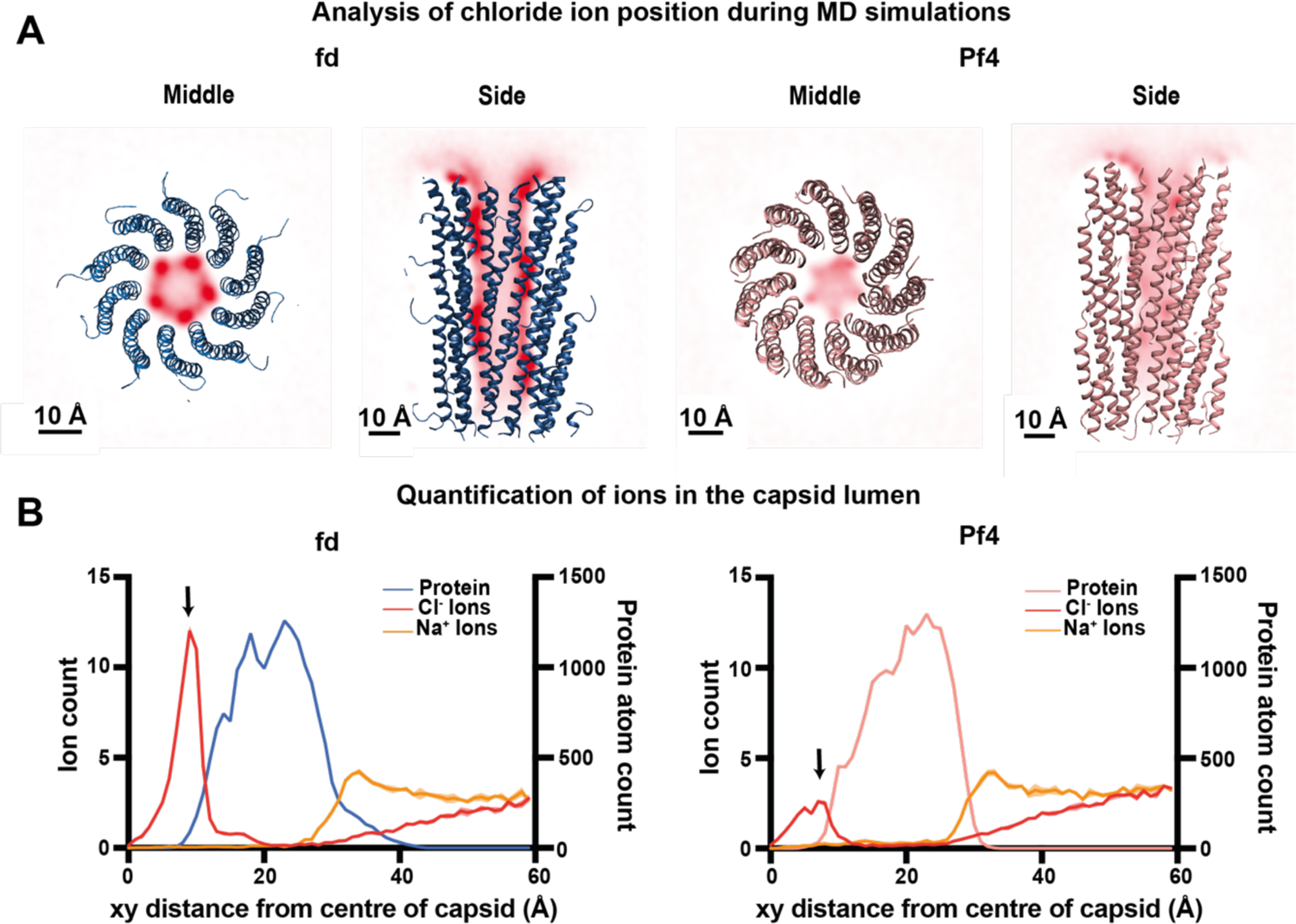
Atomistic molecular dynamics simulation analysis of fd and Pf4 capsids. **A)** Weighted ionic density of chloride ions (computed using VMD volmap) averaged over all trajectory frames. Density of chloride ions is shown in red. Dark red indicates higher ionic density. Fd capsid (left), shown in blue, depicts high levels of chloride ions in the capsid lumen (sliced at the midpoint of the filament). Pf4 capsid (right), shown in salmon, has comparatively lower levels of chloride ions in the capsid lumen (sliced at the midpoint of the filament). Both systems were simulated in 0.15 M NaCl. **B)** Quantification of ion number at different positions in a cross section of the phage, starting from the centre of the capsid for fd (left) and Pf4 (right). Fd protein atoms are coloured blue, Pf4 coloured salmon, chloride ions red and sodium ions yellow. Protein atoms are shown for clarity. The recruitment of chloride ions to the interior of the capsid can be seen as a peak at approximately 10 Å (indicated by arrows), with a higher peak observed for fd as compared to Pf4. Mean data is plotted across all four repeats (simulations) for each system in bold colour, with the standard error of the mean (SEM) plotted in transparent colour. See Figure S4E for additional analyses.

**Figure 4:**
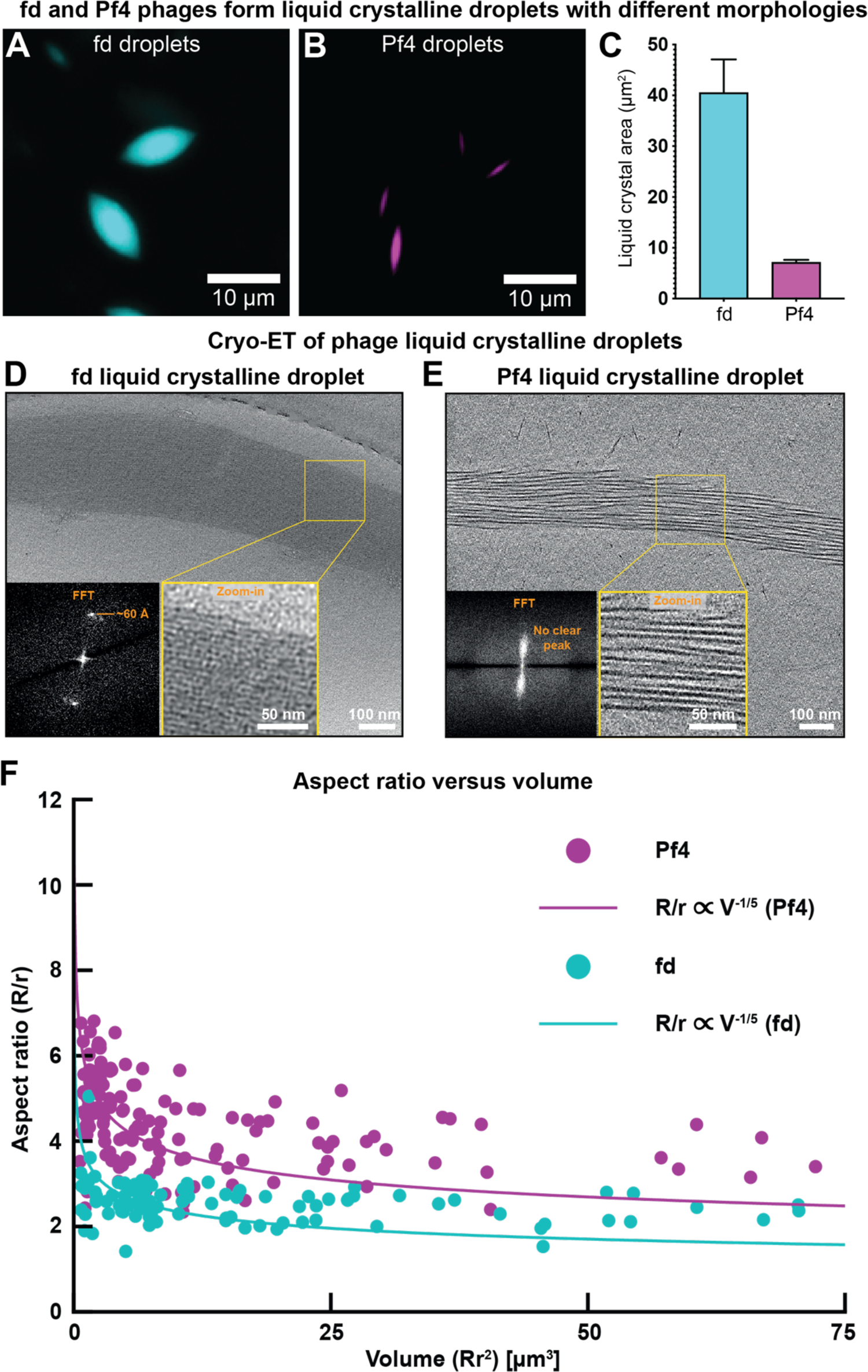
Comparison of fd and Pf4 liquid crystalline droplet morphology and filament packing. **A-B)** Representative light microscopy images of droplets formed by Alexa-488-labelled fd (cyan) and Pf4 phages (magenta). **C)** Bar chart showing average droplet area as assessed by light microscopy followed by segmentation of liquid crystalline droplets. Values shown are the mean of three independent experiments and error bars represent standard deviation. **D-E)** Droplet morphology as observed via cryo-ET. Inset: Zoom-in and Fourier Transform. **F)** Liquid crystalline droplet aspect ratio as a function of volume. Droplets, as visualised via light microscopy, follow the scaling law *R*/*r* ∝ *V*^()/’^.

### Phage geometry but not biochemistry dominates liquid crystalline droplet formation

Having resolved the structure of the fd phage capsid and described the biochemical differences with Pf4, we next investigated whether the liquid crystalline droplet-forming properties of both fd and Pf4 are different. We assembled liquid crystalline droplets by mixing phage with alginate, a biopolymer commonly found in the extracellular matrix of biofilms (Secor et al., 2015), and compared both fd and Pf4 droplets at phage and biopolymer concentrations that gave equivalent levels of total liquid crystalline droplet formation (Figures S5A-C). Under these conditions, we found that individual fd droplets have a different morphology, evidenced by a significantly higher volume and smaller aspect ratio (ratio of the major and minor elliptical axis) than individual Pf4 liquid crystalline droplets (Figures 4A-C and S5D-F). Interestingly, electron cryotomography (cryo-ET) of small liquid crystalline droplets amenable to imaging shows that fd droplets consist of laterally-packed phages, with Fourier transforms showing a regular spacing of ∼60 Å between phages. This is in contrast to liquid crystalline droplets of Pf4 showing weaker lateral ordering, but nevertheless tight overall packing, with no discrete peaks in the Fourier transforms (Figures 4D-E, Movie S2).

To understand the extent that observed liquid crystalline droplet morphologies were governed by the underlying biochemistry of the phages (Figure 2) as opposed to the phages’ geometry, we developed a physical model of liquid-crystalline droplets (also called tactoids) containing hard rods, to link the geometrical properties of the phages to those of the droplets. First, we performed a theoretical scaling calculation (Prinsen and Van Der Schoot, 2003), involving minimizing the free energy of a liquid crystalline droplet that accounts for surface (interfacial) and volumetric (elastic) contributions (see theory section in Methods for details). In agreement with previous studies (Bagnani et al., 2018, Kuhnhold and van der Schoot, 2022, Prinsen and Van Der Schoot, 2003), the scaling calculation predicts the following relationship between the geometrical properties of the droplet and its physical properties:

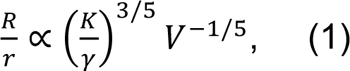

where R is the long (major) axis of the droplet, r is the short (minor) axis of the droplet, K is the Frank elastic constant, γ is the surface tension, and V is the droplet volume. The relation, which is valid for *r* ≪ *R*, predicts a decrease in the aspect ratio of the liquid crystalline droplet with an increase in the size (volume) of the droplet. As predicted, we observed that the aspect ratios and volumes of both the fd and Pf4 droplets displayed this relationship (Figure 4F). Interestingly, we found the aspect ratios of Pf4 liquid crystalline droplets to be larger than the aspect ratios of the fd droplets at similar volumes (Figure 4F).

To explain the observed difference between the Pf4 and fd curves in Figure 4F, we performed a second scaling calculation to link the physical droplet properties in the prefactor of Eq. (1) to the phage geometry (see Methods). By approximating the elastic constant K and the surface tension γ, we derived the following scaling relationship:

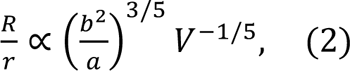

where b is the length (major axis) of the phage (fd 0.9 μm and Pf4 3.8 μm), and a is the width (minor axis) of the phage (fd 64 Å and Pf4 62 Å). The above relation, which contains only geometrical properties of the phages and droplets, predicts that for droplets of similar volume, a larger phage length *b* should correspond to a larger droplet aspect ratio. Both fd and Pf4 phages have similar widths, but Pf4 phages are longer; as predicted, this is in line with the trend for larger aspect ratios of Pf4 droplets than fd droplets at similar volumes (Figure 4F). Biochemical phage interactions could affect the physical prefactors in Eq. (1), and thus could enter Eq. (2), but do not seem to underlie the qualitative differences in our measurements of droplet aspect ratios. This suggests that our biophysical model of tactoids containing hard rods is sufficient to explain the observed differences between Pf4 and fd liquid droplets in Figure 4F. Taken together, our experimental measurements and predictions from the scaling theory suggest that overall droplet morphology is governed by biophysical effects through droplet size and phage geometry.

### Droplet encapsulation and antibiotic protection of bacterial cells

Since biophysical effects, rather than biochemical properties of phages, appear to govern droplet morphology, we next asked how these morphologically diverse droplets interact with rod-shaped *E. coli* and *P. aeruginosa* bacteria. Previous studies have shown that liquid crystalline droplets formed by Pf4 can fully encapsulate *P. aeruginosa* cells, which correlate with protection from antibiotic treatment (Tarafder et al., 2020). To assess what drives encapsulation of cells by liquid crystalline droplets, we imaged fd and Pf4 droplets in the presence of *E. coli* and *P. aeruginosa* cells, respectively representing their native host bacterium as well as a non-host bacterium. Fd droplets were associated with both *P. aeruginosa* (non-host) and *E. coli* (native host) cells (Figures 5A and 5D). Similarly, Pf4 droplets could associate with *P. aeruginosa* cells (native host), as described previously, as well as *E. coli* (non-host) (Figures 5B and 5E), suggesting that bacterial cell association is a biophysical process rather than being driven by specific biochemical interactions, since the surfaces of the phages as well as the outer surfaces of *E. coli* and *P. aeruginosa* are biochemically distinct.

**Figure 5:**
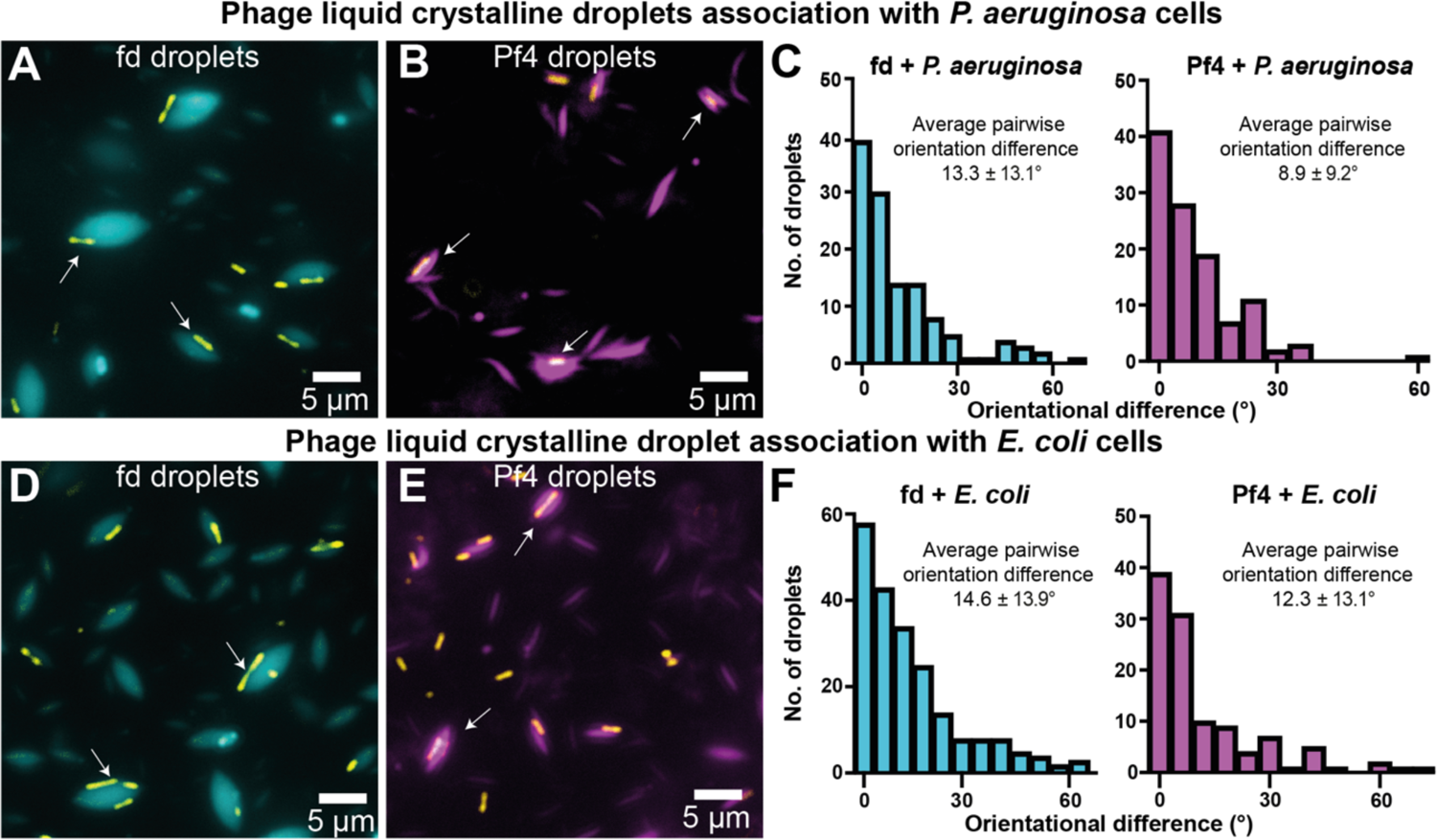
Comparison of fd and Pf4 liquid crystalline droplet association with bacterial cells. **A-B)** Representative images showing association of fd and Pf4 liquid crystalline droplets with *P. aeruginosa* cells observed in light microscopy. Transmitted light channel shows bacteria (yellow) and fluorescence channel shows fd (cyan) and Pf4 (magenta) droplets, respectively. **C)** Histogram of pairwise orientational differences between bacterial cells and associated liquid crystalline droplets from semi-automated segmentation of images. Values reported are the mean and standard deviation from three independent experiments. Left panel fd + *P. aeruginosa* (n=144), right panel Pf4 + *P. aeruginosa* (n=163), significantly different with *** P_value_ < 0.001. **D-E)** Corresponding representative images showing association of fd and Pf4 droplets with *E. coli*. Transmitted light channel shows bacteria (yellow) and fluorescence channel shows fd (cyan) and Pf4 (magenta) droplets, respectively. **F)** Histogram of pairwise orientational differences between bacterial cells and associated droplets from semi-automated segmentation of images. Values reported are the mean and standard deviation from three independent experiments. Left panel: fd + *E. coli* (n=244), right panel: Pf4 + *E. coli* (n=137), no significant difference. Pf4 + *P. aeruginosa* versus Pf4 + *E. coli*, * P_value_ < 0.05. Fd + *P. aeruginosa* versus fd + *E. coli*, no significant difference. Pf4 + *P. aeruginosa* versus fd + *E. coli*, *** P_value_ < 0.0001. Fd + *P. aeruginosa* versus Pf4 + *E. coli*, no significant difference.

Curiously, although both phage droplets could associate with bacterial cells, Pf4 droplets more readily encapsulated both bacteria and were able to completely surround the cell, whereas fd droplets interacted with cells laterally (Figures 5A-F). Using semi-automated analysis of multiple droplet:cell images, we measured the pairwise orientation differences between the long axes of bacterial cells and their associated droplets (Figures 5C and 5F). Our measurements show an alignment of axes in all cases, with the tightest alignment between Pf4 droplets and *P. aeruginosa* (8.9 ± 9.2°; significantly different from Pf4 with *E. coli* 12.3 ± 13.1°, P_value_ < 0.001). The orientational difference of fd droplets with both bacteria was larger, but still alignment was observed, suggesting the interaction is not stochastic (fd with *P. aeruginosa* 13.3 ± 13.1°, fd with *E. coli* 14.6 ± 13.9°; no statistically significant difference) (Figures 5C and 5F). Fd droplets associated with bacteria were also larger than Pf4 droplets associated with both bacteria tested, with fd droplets ∼2.5-2.9 μm larger compared to Pf4 droplets ∼1.2-1.4 μm larger than the bacterial cell major axis (Figures S6A-D). This suggests that Pf4 droplets are ideally matched with *P. aeruginosa* or *E. coli* bacterial cells, allowing encapsulation of cells, whereas the larger fd droplets tend to interact laterally with cells rather than tightly encapsulating them (Figure S6E-F).

In our previous work, we showed that encapsulation was tightly linked with antibiotic protection (Tarafder et al., 2020). To test how differences in cellular association with droplets affected antibiotic protection, we performed a previously developed antibiotic protection assay (Tarafder et al., 2020) that measures bacterial survival in different conditions against an antibiotic challenge. Our assay showed that despite clear and quantifiable differences between fd and Pf4 droplet association with cells (Figures 5 and S6), both fd and Pf4 droplets could protect *P. aeruginosa –* corroborating previous reports (Figure 6A, (Tarafder et al., 2020, Secor et al., 2015)). Surprisingly, we found that these droplets could even protect *E. coli* bacteria from antibiotic challenge (Figure 6B), which shows that association with liquid crystalline droplets is sufficient for a bacterium to survive antibiotic treatment. These antibiotic protection assays suggest that biophysical parameters govern and profoundly affect bacterial responses to external stresses such as antibiotic treatment.

**Figure 6:**
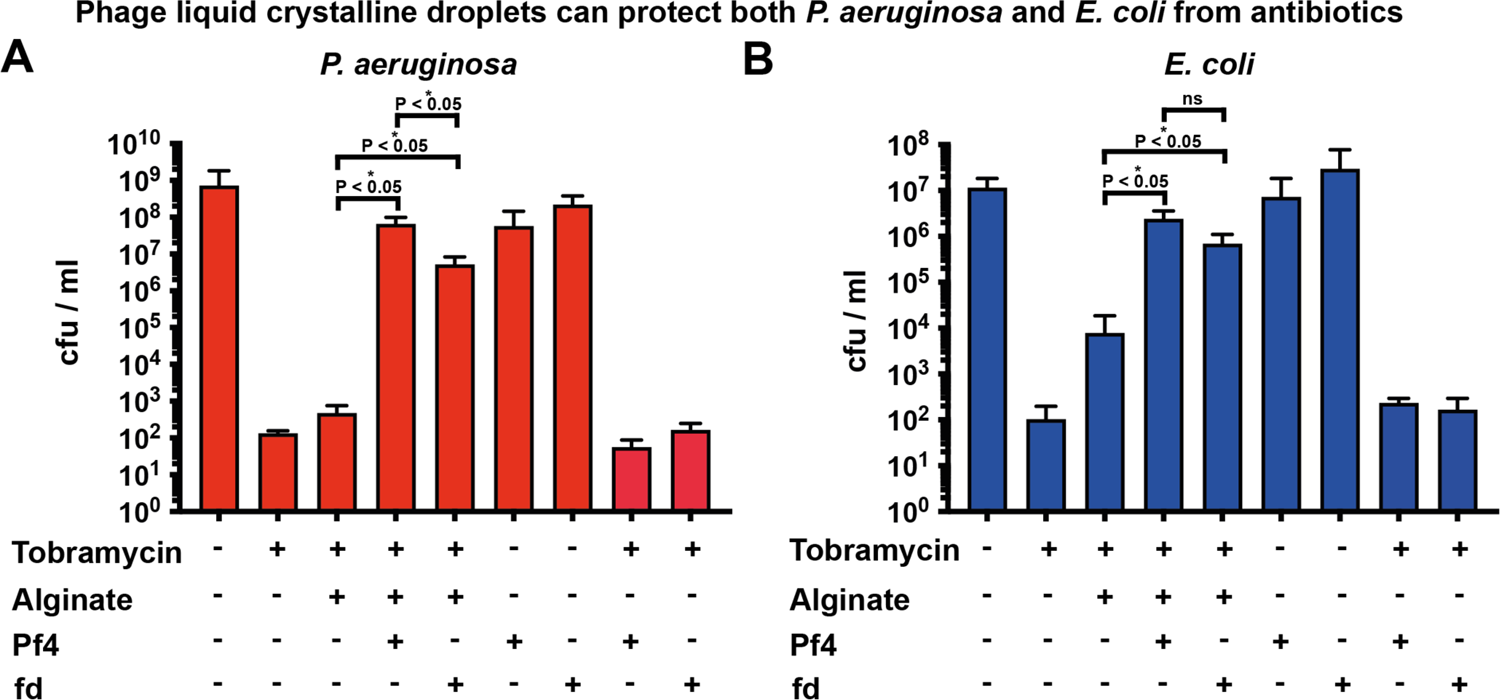
Antibiotic protection assay. Bar graphs showing viability of cells in colony-forming units (cfu) per ml (y-axis) in the presence of different reagents (x-axis) from three independent experiments for **A**) *P. aeruginosa* and **B**) *E. coli* against tobramycin treatment. Both Pf4 and fd liquid crystalline droplets protect *P. aeruginosa* to a significantly greater extent than sodium alginate alone (P_value_ < 0.05). Fd droplets protected significantly less well than Pf4 droplets (P_value_ < 0.05). Both Pf4 and fd droplets protect *E. coli* to a significantly greater extent than sodium alginate alone (P_value_ < 0.05). No significant difference (n.s.) was observed between the level of protection provided by fd and Pf4 droplets.

## Discussion

In this study, we have solved the capsid structure of an archetypal class I inoviral bacteriophage, fd, which has been intensely studied, being the first discovered inovirus, used in biotechnology for phage display (Smith, 1985) as well as for investigating phase transitions of colloidal rod-like molecules (Dogic and Fraden, 2001). Numerous inconsistent models for the fd phage capsid have been proposed over the years by X-ray fibre diffraction, ssNMR and 8 Å-resolution cryo-EM (Marvin, 1966, Cross and Opella, 1985, Hunter et al., 1987, Colnago et al., 1987, Marvin et al., 1994, Zeri et al., 2003, Marvin et al., 2006, Wang et al., 2006). Our 3.2 Å-resolution cryo-EM structure settles this long-standing debate and shows how individual capsid monomers form a cylindrical phage with five-fold symmetry (Figure 1). Also notably, previous studies had suggested that the Y21 residue within the pVIII capsid protein induces structural flexibility, and that the mutation Y21M could reduce this flexibility (Tan et al., 1999). Our structure suggests that while Y21 is involved in hydrophobic interactions within the capsid, it does not act as a structural hinge, evidenced by our high-resolution map. While previous cryo-EM studies had suggested there may be a structural continuum of fd structures (Wang et al., 2006), we did not capture these states in our data set.

Comparing the class I fd capsid structure with the class II Pf4 phage reveals four lysine residues extending into the lumen (compared with two positively charged residues in Pf4), where they could interact closely with ssDNA (Figure 2). While density for DNA could be detected in our map (Figure S1F), it did not allow for unambiguous atomic model building, consistent with previous studies on the Ike phage (Figure S2). The size of the fd phage lumen, its increased positive charge, along with our MD simulations (Figure 3), suggest that the DNA arrangement in this phage is markedly different to Pf4. Previous literature suggested that fd contains a circular single-stranded DNA genome (Marvin, 1966), and this is supported by our data as the linear ssDNA in Pf4 would require less positively charged residues for encapsidation. Given that fd has a comparable number of capsid proteins per length unit, the fact that the fd capsid can compensate twice as many negative charges as Pf4 suggests that its DNA genome indeed is circular.

Fd and Pf4 form liquid crystalline droplets with starkly differing morphology; at the same overall specimen droplet volumes, individual fd droplets had a smaller aspect ratio and significantly higher volume than Pf4 droplets, which were smaller and elongated (Figure 4). Even though the underlying biochemistry and surface charge distributions of both phages are distinct, these have minimal influence on droplet morphology. According to our theoretical calculations and predictions, differences in the size and shape of constituent phages are sufficient to explain differences in emergent droplet morphology. This result shows that biophysical characteristics, rather than biochemical, dominate in the formation of droplets. Similar biophysical governance was also observed in droplet association with cells (Figure 5): Pf4 droplets almost fully encapsulate rod-shaped *P. aeruginosa* and *E. coli* cells, compared to the artificial case of fd droplets, where only lateral interactions were seen. Combined with previous studies (Secor et al., 2015, Tarafder et al., 2020), this suggests the intriguing possibility that shape complementarity of the Pf4 droplets to the *P. aeruginosa* cell could have been evolutionarily selected, e.g. by alteration of phage length, to maximise bacterial protection and thus fitness in harsh environments.

One of the most striking results in this study was that droplet association with cells is sufficient for antibiotic protection (Figure 6), rather than complete encapsulation. Even though fd droplets did not shape-complement cells, their presence nonetheless provided protection against antibiotic treatment to rod-shaped bacteria (Figure 6). It is tempting to speculate that the viscous surrounding created by droplets leads to reduced antibiotic diffusion and reduced access to the bacterial cell, conferring protection. However, further experiments will be needed to test this, and detailed mechanisms of how droplets mediate protection against antibiotic treatment will be an exciting future direction of experimental and theoretical research. Since bacteria often proliferate in environments rich in rod-like or filamentous molecules accompanied by biopolymers, for example in the biofilm matrix (Tarafder et al., 2020, Böhning et al., 2022, Secor et al., 2015) or on tissues covered with mucus (Wagner et al., 2018), our structural, biochemical, and imaging data suggest a paradigm where the biophysics of such associations strongly influence bacterial interactions with, and response to, external stimuli (Figure 7).

**Figure 7:**
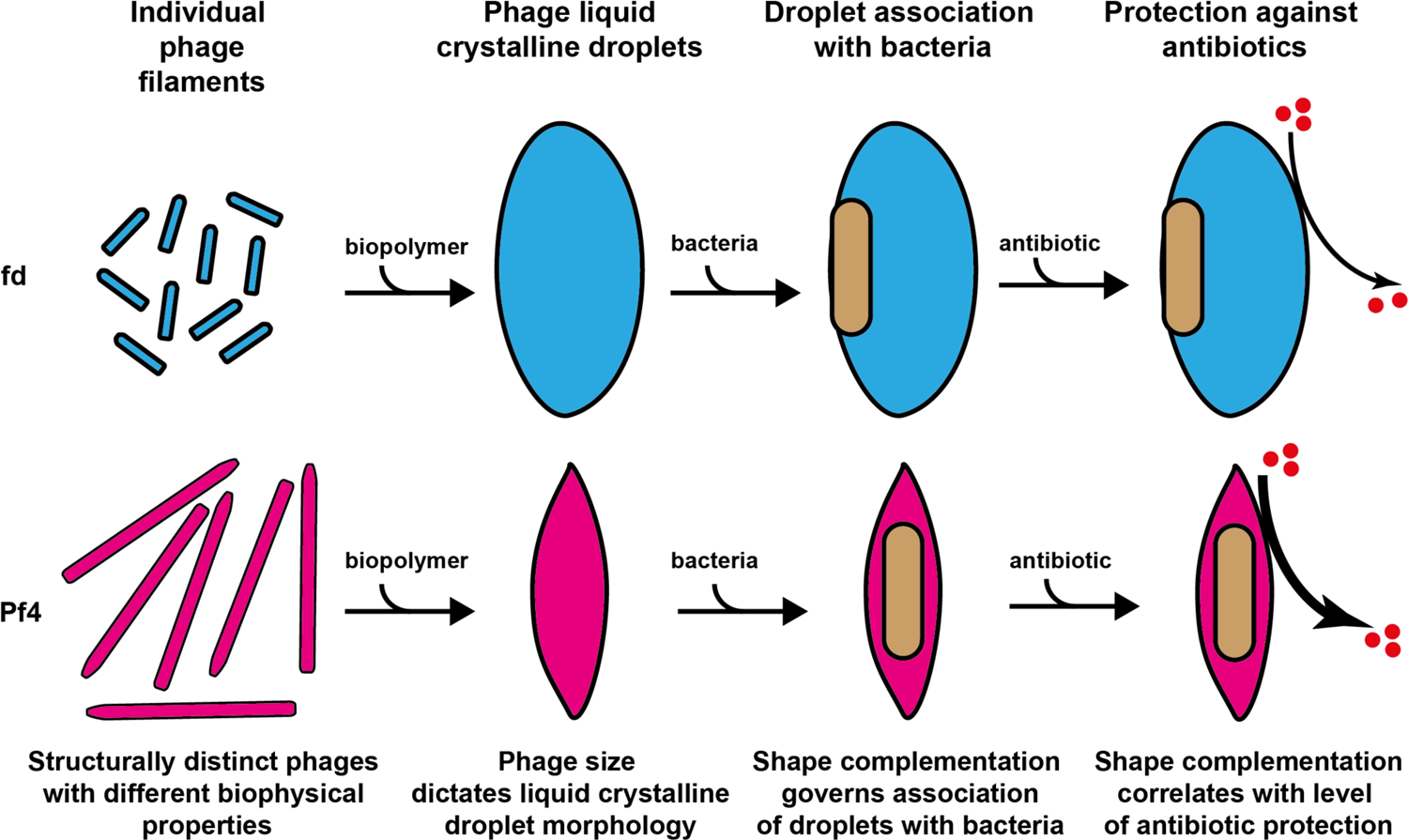
Schematic depicting biophysical nature of phage liquid crystalline droplet-mediated antibiotic tolerance of bacteria. The structurally and biochemically distinct inoviruses, fd and Pf4, form liquid crystalline droplets in the presence of biopolymer with diverse morphologies dictated by their biophysical size, rather than biochemical characteristics. Both fd and Pf4 phage liquid crystalline droplets associate with bacterial cells, with the level of association dictated by biophysical size complementation of the droplet with the rod-shaped bacteria. Phage liquid crystalline droplet association with bacteria may form a diffusion barrier that results in increased antibiotic tolerance, which correlates with the level of bacterial cell encapsulation by the droplet.

## Methods

### Bacterial strains and growth conditions

*E. coli* strains ER2738 (a kind gift from Prof. Eric Grelet, Centre de Recherche Paul Pascal) and XL1 were used for fd phage amplification and encapsulation assays respectively. *P. aeruginosa* strains PAO1 or PAO1 *ΔPA0728* (a kind gift from Prof. Patrick Secor, University of Montana) were used for experiments as indicated. Unless otherwise specified, shaking cultures of bacteria were incubated at 37°C in Luria-Bertani (LB) medium with agitation at 180 rpm (revolutions per minute).

### Phage production and purification

Fd was amplified from a starter preparation of purified fd (a kind gift from Prof. Eric Grelet, Centre de Recherche Paul Pascal) using ER2738 *E. coli*. Briefly, to propagate fd phage, a 250 ml culture of ER2738 *E. coli* at OD_600_ 0.3-0.4 was infected with 100 μl of fd phage at 1 mg/ml and grown overnight at 37°C with agitation. Cells were pelleted by centrifugation (5,000 *g*, 20 minutes, 4 °C) and the supernatant further centrifuged to remove residual cells (15,000 *g*, 30 minutes, 4 °C). The supernatant was adjusted to 0.5 M NaCl and phage precipitated overnight with 10% (w/v) PEG (polyethylene glycol) 8000. Precipitated phage was harvested by centrifugation (12,000 *g*, 30 minutes, 4 °C). The phage containing pellet was resuspended in PBS (phosphate buffered saline) and dialysed against PBS overnight using 10 kDa MWCO (molecular weight cut-off) snakeskin dialysis membranes (ThermoFisher). Pf4 was isolated from static PAO1 biofilms as described previously (Tarafder et al., 2020). For further amplification of Pf4, 1 x 10^3^ pfu (plaque forming units)/ml of the Pf4 isolated from PAO1 biofilms were incubated with 1 ml of PAO1 culture at 0.5 OD_600_ for 15 minutes and mixed with hand-hot 0.8% (w/v) agar. The mix was plated onto 10 cm^2^ LB-agar plates and incubated overnight at 37 °C. Each plate was covered with 5 ml PBS and incubated for 6 hours before the PBS was collected, centrifuged (12,000 *g*, 30 minutes, 4°C) and the supernatant adjusted to 0.5 M NaCl and phage precipitated with 10% (w/v) PEG 6000. Precipitated phage was harvested by centrifugation (12,000 *g*, 30 minutes, 4 °C). The phage containing pellet was resuspended in PBS and dialysed against PBS overnight using 10 kDa MWCO snakeskin dialysis membranes (ThermoFisher). Yield of both fd and Pf4 phage preparations was estimated using Nanodrop (Thermo Scientific).

### Fluorescent labelling of phage

Purified fd or Pf4 phage was dialysed into 10 mM sodium carbonate buffer pH 9.2 using a 10 kDa MWCO snakeskin dialysis membrane (ThermoFisher). One ml of fd or Pf4 phage (5 mg/ml) was incubated with 100 µg A488 fluorescent dye (ThermoFisher) for 1 hour at room temperature (RT) with end-over-end agitation. A488-Labelled phage was isolated from free dye by passing over two PD10 desalting columns (GE Healthcare).

### Cryo-EM grid preparation

Cryo-EM grids were prepared by pipetting 2.5 µl of sample onto glow-discharged Quantifoil grids (Cu/Rh R2/2, 200 mesh) and plunge frozen into liquid ethane using a Vitrobot Mark IV (ThermoFisher). For cryo-ET only, 7 µl of sample was mixed with 1 µl of protein-A conjugated with 10 nm colloidal gold (CMC, Utrecht) before 2.5 µl was applied to the grid. Plunge-frozen grids were transferred to and stored in liquid nitrogen until imaging.

### Cryo-EM and cryo-ET data collection

Cryo-EM data for screening specimens were collected using a Talos Arctica (ThermoFisher) operated at 200 kV. High-throughput data was collected on a Titan Krios microscope operated at 300kV fitted with a Quantum energy filter (slit width 20 eV) and a K3 direct electron director (Gatan) operating in counting mode at an unbinned, calibrated pixel size of 1.1 Å using the EPU software. A combined total dose of approximately 53.9 e^-^/A^2^ per exposure was applied with each exposure lasting 3.6 s and 40 frames were recorded per movie. In total 4259 movies were collected between −1 to −3 µm defocus. Tilt series data for cryo-ET was collected on a Titan Krios using the Quantum energy filter and K2 direct electron director with SerialEM software (Mastronarde, 2005). Tilt series were collected in two directions starting from 0° between ±60° with a 1° tilt increment, acquired with defoci ranging from −4 to −5 µm, with a combined dose of approximately 120 e^-^/Å^2^ applied over the entire series. Tilt series were collected at an unbinned calibrated pixel size 4.41 Å.

### Cryo-EM data processing

Helical reconstruction was performed in RELION 3.1 (Scheres, 2012, He and Scheres, 2017, Zivanov et al., 2018, Zivanov et al., 2020). Movies were motion-corrected using the RELION implementation of MotionCor2 (Zheng et al., 2017), and defocus was estimated using CtfFind4 (Rohou and Grigorieff, 2015). Using a previously established starting symmetry of fd of ∼36° and ∼16 Å (Marvin et al., 1994, Wang et al., 2006), iterative rounds of 3D refinement and 3D classification were used to create a reference with alpha-helical features that enabled refinement to <5 Å resolution. CTF refinement and Bayesian polishing with CTF-multiplication were then applied to create a 3.2 Å resolution map. Symmetry searches were used during classification and refinement, and a final rotation of 35.46° and rise of 16.62 Å per subunit were obtained.

### Theoretical scaling calculation of liquid crystalline droplet geometry

We performed a theoretical scaling calculation, involving minimizing an approximate free energy *F* of a liquid crystalline droplet that comprises of hard rods (modelled phages) (Prinsen and Van Der Schoot, 2003). By assuming that the volume of a tactoid is much greater than the volume of an individual rod, we can decompose the total droplet free energy into two terms: one term describing the surface (interfacial) contribution and a second term describing the volumetric (elastic) contribution.

In order to arrive at a free energy that is simple enough to solve, yet complex enough to retain information on the overall geometry of the rods and droplets, we make further physical assumptions. First, we use what is called the one-constant approximation for the elastic term (de Gennes and Prost, 1993). This means that the splay and bend elastic effects contribute to the free energy in equal measure. Second, we ignore twist and saddle-splay effects, as we are primarily concerned with large changes in overall shape and size, not on complex internal organization of the rods. Lastly, we assume that the droplet is sufficiently elongated (not spherical) and that there is a slight curvature in the stacking of rods in the droplet, as observed in our experiments. Given the above assumptions, the free energy is given by

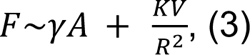

where γ is the surface tension, *A(r,R)* is the area of the droplet, *V(r,R)* is the droplet volume, *K* is the Franck elastic constant, *r is the short (minor)* axis of the droplet, and *R* is the long (major) axis of the droplet.

To find the equilibrium geometry of the droplet, we minimize *F* with respect to both *r* and *R* at fixed volume. In doing so we approximate the area as *A*∼*Rr* and the volume as *V*∼*Rr*^-^. The equations to satisfy are given by:

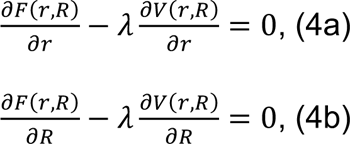

where λ is a Lagrange multiplier. By solving the above equations, rearranging them for λ, and then setting them to be equal, we obtain the following relation:

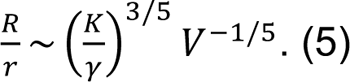

Next, to involve the shapes of the hard rods we approximate the Frank elastic constant and interfacial tension with their values at the homogeneous-bipolar transition given by (van der Schoot, 1999):

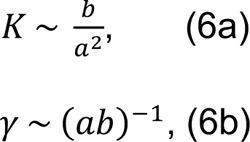

where we have assumed a constant volume fraction of hard rods in a droplet. Given this, we reach the final relation given by:

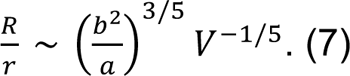

### Antibiotic protection assay

An overnight culture of PAO1 *ΔPA0728* was grown in LB media at 37 °C, diluted 1 in 100 into LB medium and grown at 37 °C to an OD_600_ of 0.5. Hundred µl of the resulting culture was added to a 96-well plate and grown for a further 30 minutes at 37 °C. Hundred µl of the indicated phage and/or polymer components were added to the culture such that final concentrations of components were: sodium alginate (Scientific Laboratory Supplies) (4 mg/ml), fd (1 mg/ml) and Pf4 (1 mg/ml). Additionally, tobramycin (10 µg/ml) (Sigma) was added as indicated and cultures grown further for 3 hours before a 10 µl sample for each assay condition was taken, serially diluted 10-fold and 100 µl of the dilutions plated onto LB agar plates. Plates were incubated overnight at 37 °C and colonies forming units (cfu) enumerated. Experiments were performed in triplicate. Mean cfu/ml with standard deviation were calculated and plotted using Prism GraphPad software.

### Fluorescence microscopy

#### Phage liquid crystalline droplets

To obtain equivalent levels of total liquid crystalline droplet formation (summed area of all liquid crystalline droplets), A488-labelled fd phage (final concentration 0.5 mg/ml) or A488-labelled Pf4 phage (final concentration 2.7 mg/ml) were mixed with sodium alginate (final concentration 4 mg/ml) and incubated at room temperature for 24 hours. Five µl of the resulting sample was pipetted onto 0.7% (w/v) agar pads constructed using 15 x 16 mm Gene Frames (ThermoFisher) following the manufacturer’s protocol, with a coverslip placed on top. The slide was imaged using a Zeiss Axioimager M2 (Carl Zeiss) microscope in both brightfield and fluorescence mode. Quantification of individual liquid crystalline droplet area and morphology was performed using MicrobeJ (Ducret et al., 2016). Maxima corresponding to individual liquid crystalline droplets were analysed using MicrobeJ shape analysis functions to quantify droplet area, major axis length and minor axis length. Experiments were performed in triplicate. Presented images were background subtracted and figure panels prepared using Fiji. Graphs were plotted using Prism GraphPad software.

#### Phage liquid crystalline droplets / bacterial cells

*P. aeruginosa* PAO1 *ΔPA0728* or *E. coli* XL1 were grown to an OD_600_ of 0.5 and incubated with A488-labelled phage (final concentration 1 mg/ml) and sodium alginate (final concentration 4 mg/ml) for 3 hours. Five µl of the sample was pipetted onto 0.7% (w/v) agar pads constructed using 15 x 16 mm Gene Frames (ThermoFisher) with a coverslip applied, and imaged using a Zeiss Axioimager M2 microscope (Carl Zeiss). Images were background subtracted and figure panels prepared using Fiji.

#### Semi-automated segmentation of images with phage liquid crystalline droplets surrounding bacterial cells

Bacteria were manually selected from brightfield channel images to identify the positions of cells. Bacterial cell shapes were found using the activecontour algorithm in Matlab (Chan and Vese, 2001). The regions of identified bacteria were dilated and used as seed inputs for the segmentation of the liquid crystalline droplets in the fluorescence channel using the activecontour algorithm. The segmented bacterial cells from the bright field and the liquid crystalline droplets from the fluorescence channel were then used to calculate morphological parameters.

#### Preparation of molecular dynamics simulation systems

The previously published Pf4 capsid cryo-EM structure (PDB: 6TUQ) (Tarafder et al., 2020) and the fd capsid cryo-EM structure described in this paper were cropped using VMD (Humphrey et al., 1996) and PyMOL (Schrodinger, 2015) to prepare simulations of a smaller and computationally judicious system. The Pf4 and fd capsid systems were centred in a 127.3 Å^3^ and 126.2 Å^3^ box respectively. Both systems were solvated using the TIP3P water model (Jorgensen et al., 1983) and neutralised in 0.15 M NaCl. One round of steepest descent energy minimisation was performed on each system for 100 ps, followed by one 5 ns NVT and one 5 ns NPT equilibration, with restraints applied to the heavy backbone atoms.

#### Atomistic molecular dynamics simulations

Simulations were performed using the Gromacs 2021.3 simulation package (Abraham et al. (2015), https://doi.org/10.5281/zenodo.5053220) and the CHARMM-36 forcefield (Best et al., 2012). Atomistic simulations were run in quadruplicate for 250 ns using a 2 fs timestep. Periodic boundary conditions were applied and the velocity-rescale thermostat (Bussi et al., 2007), with a coupling constant of 0.2 ps, was used to maintain temperature at 310 K. Pressure was maintained at 1 bar using the Parrinello-Rahman barostat (Parrinello and Rahman, 1981) with a coupling constant of T_p_ = 2.0 ps and compressibility of 4.5 x10^-5^ bar^-1^. Long range electrostatic interactions were modelled using the Particle-Mesh Ewald (PME) method (Darden et al., 1993) and a 1.2 nm cut-off was used. Van der Waals interactions were switched between 1.0 and 1.2 nm using the force-shift modifier. Dispersion correction was not applied. The LINCS algorithm (Hess et al., 1997) was used to constrain bonds to their equilibrium values. During both Pf4 and fd capsid simulation production runs, 50 kJ/mol/nm^2^ restraints were applied in xyz on the bottom and top c-alpha carbons of the protein (z < 3 or z > 9). See Figure S4A-B for views of restrained atoms and simulation RMSD values. Analysis was performed using a PyLipID script (Song et al., 2022), with modified cut-offs (0.3,0.5) for ion binding, gmx tools (Abraham et al. (2015), https://doi.org/10.5281/zenodo.5053220) and VMD volmap (Humphrey et al., 1996).

For all volmap and PyLipID analyses, trajectories sampled every five frames from each repeat were used for the analysis, totalling 20,000 frames per capsid simulation system. For PyLipID analysis, the trajectories were further sampled by 10 frames for Pf4 and 40 frames for fd. For quantification of ions within the lumen, the cylayer command within MDAnalysis (Michaud-Agrawal et al., 2011, Gowers et al., 2016) was used to count ions within cylindric layers of 1 Å intervals positioned on the centre of geometry of the capsid atoms. Analysis was conducted every 5 ns from the full 250 ns trajectories to a maximum distance of 60 Å. Graphs were plotted using Plotly (Plotly Inc.) and Prism GraphPad software. Preparation of systems for simulation was performed using computing resources at The Kavli Institute in the Structural Bioinformatics and Computational Biochemistry unit at the University of Oxford. Production runs were simulated for 250 ns using computing resources on ARCHER2.

#### Statistical analysis

Statistical analysis was performed using Prism GraphPad software and an unpaired t-test was used to calculate p-values.

## Acknowledgments

T. A. M. B. would like to thank UKRI MRC (Programme MC_UP_1201/31), the Human Frontier Science Programme (Grant RGY0074/2021), the Vallee Research Foundation, the European Molecular Biology Organization, the Leverhulme Trust and the Lister Institute for Preventative Medicine for support. P. P. is supported by a UKRI Future Leaders Fellowship [MR/V022385/1]. This project made use of time on ARCHER2 granted via the UK High-End Computing Consortium for Biomolecular Simulation, HECBioSim (www.hecbiosim.ac.uk), supported by EPSRC (grant no. EP/R029407/1). We would like to thank Professor Mark Sansom and The Kavli Institute Structural Bioinformatics and Computational Biochemistry unit at the University of Oxford for facilitating the Molecular Dynamics performed in this research. We thank Jeanne Salje and Paul Edelstein for critical comments and the MRC-LMB cryo-EM facility for technical support.

## Author contributions

Conceptualisation: J.B., A.K.T., T.A.M.B.; Methodology: J.B., L.K.D., P.P., A.K.T., T.A.M.B.; Formal Analysis: J.B., M.G., S.C.L., L.K.D., U.S., R.A.C., P.J.S., P.P., A.K.T., T.A.M.B.; Investigation: J.B., M.G., S.C.L., L.K.D., P.P., A.K.T., T.A.M.B.; Writing – Original Draft: J.B., A.K.T., T.A.M.B.; Writing – Review & Editing: J.B., M.G., S.C.L., L.K.D., U.S., R.A.C., P.J.S., P.P., A.K.T., T.A.M.B.; Supervision: R.A.C., P.J.S., P.P., A.K.T., T.A.M.B; Funding Acquisition: P.J.S., P.P., T.A.M.B.

## Declaration of Interests

The authors declare no competing interests.

## Supplementary Figures

**Figure S1:**
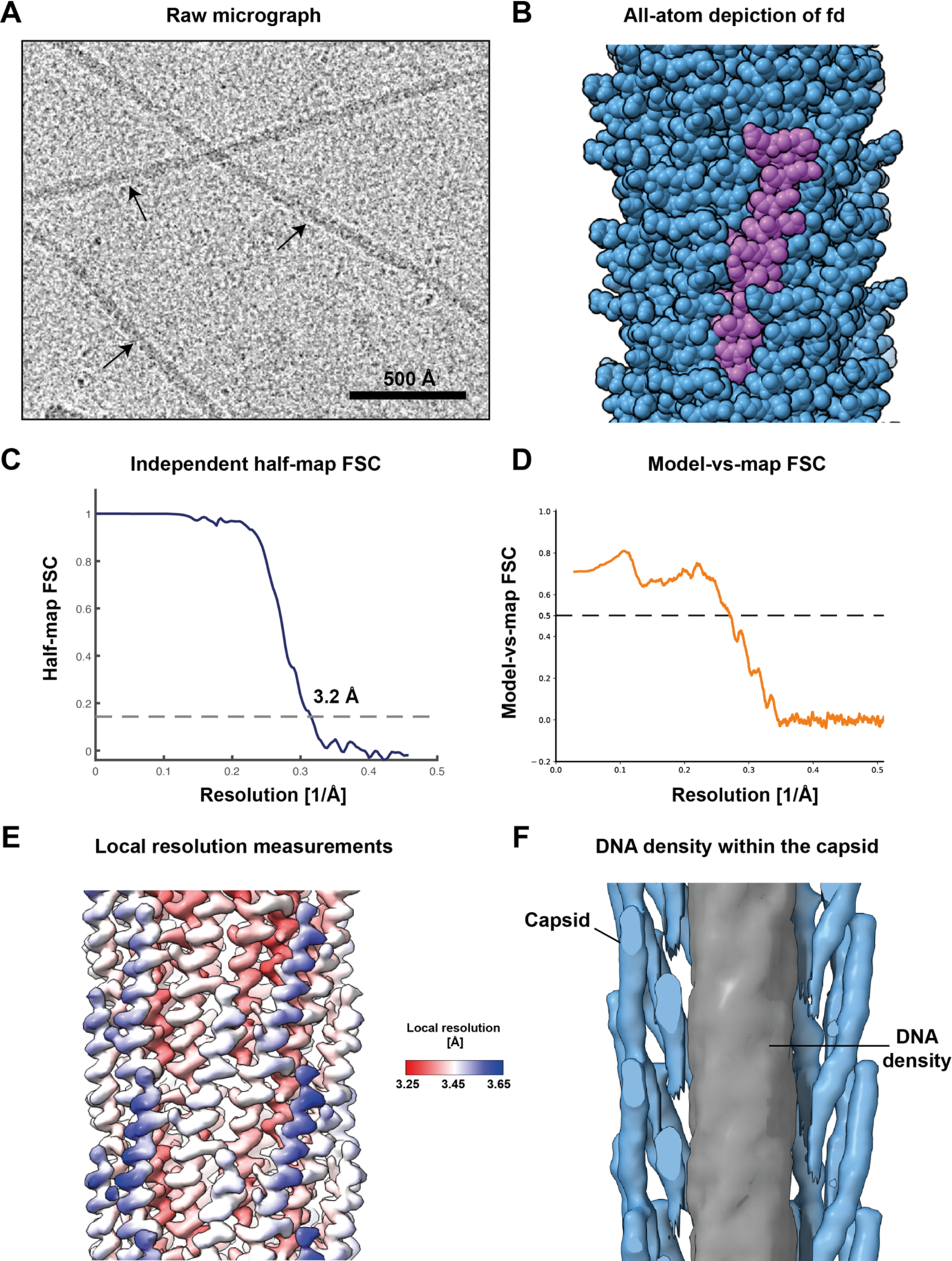
Cryo-EM structure of the bacteriophage fd at 3.2 Å resolution. **A)** Cryo-EM image of native fd phage specimen used for structure determination. **B)** All-atom sphere depiction of the bacteriophage as shown in Figure 1A, with one capsid subunit marked in purple. **C)** Fourier Shell Correlation (FSC) as calculated by RELION. The 0.143 criterion used for resolution estimation is shown as a dotted line. **D**) Model vs map FSC as calculated by PHENIX. **E)** Local resolution measurement of the cryo-EM structure. Slight differences in local resolution between individual capsid proteins indicate minor structural flexibility within the capsid. **F)** Sliced side view of cryo-EM density refined with C1 symmetry (at lower resolution). Density (2*σ* contour level) corresponding to capsid (blue) and DNA (grey) are highlighted.

**Figure S2:**
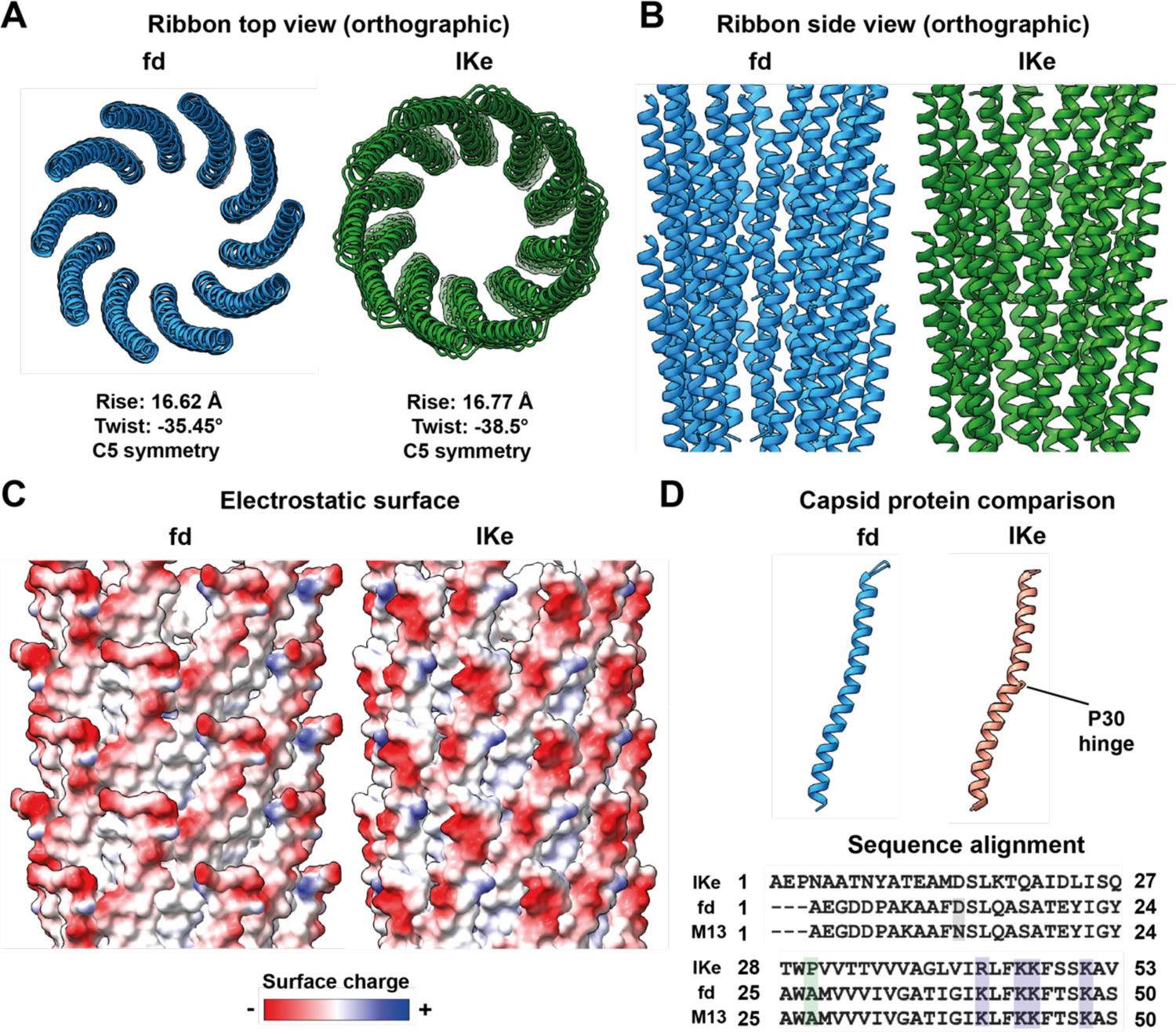
Comparison of fd with another class I bacteriophage IKe. **A)** Orthographic top view of fd versus IKe, showing a higher twist in the case of IKe. **B)** Side view of fd and IKe capsids, excluding their N-terminal residues that are not resolved in the cryo-EM density. **C)** Comparison of electrostatic surface of full-length fd and IKe reveals a negatively charged capsid surface in both cases, with three negatively charged residues in the disordered N-terminus in the case of fd. **D)** While showing a similar overall morphology, a proline residue (P30) induces a hinge in the pVIII protein of IKe. A Clustal Omega sequence alignment of the major capsid proteins of IKe, fd and M13 are shown. IKe P30 is highlighted in green and positively charged residues at the C-terminus highlighted in purple. The residue differing between fd and M13 is marked in grey.

**Figure S3:**
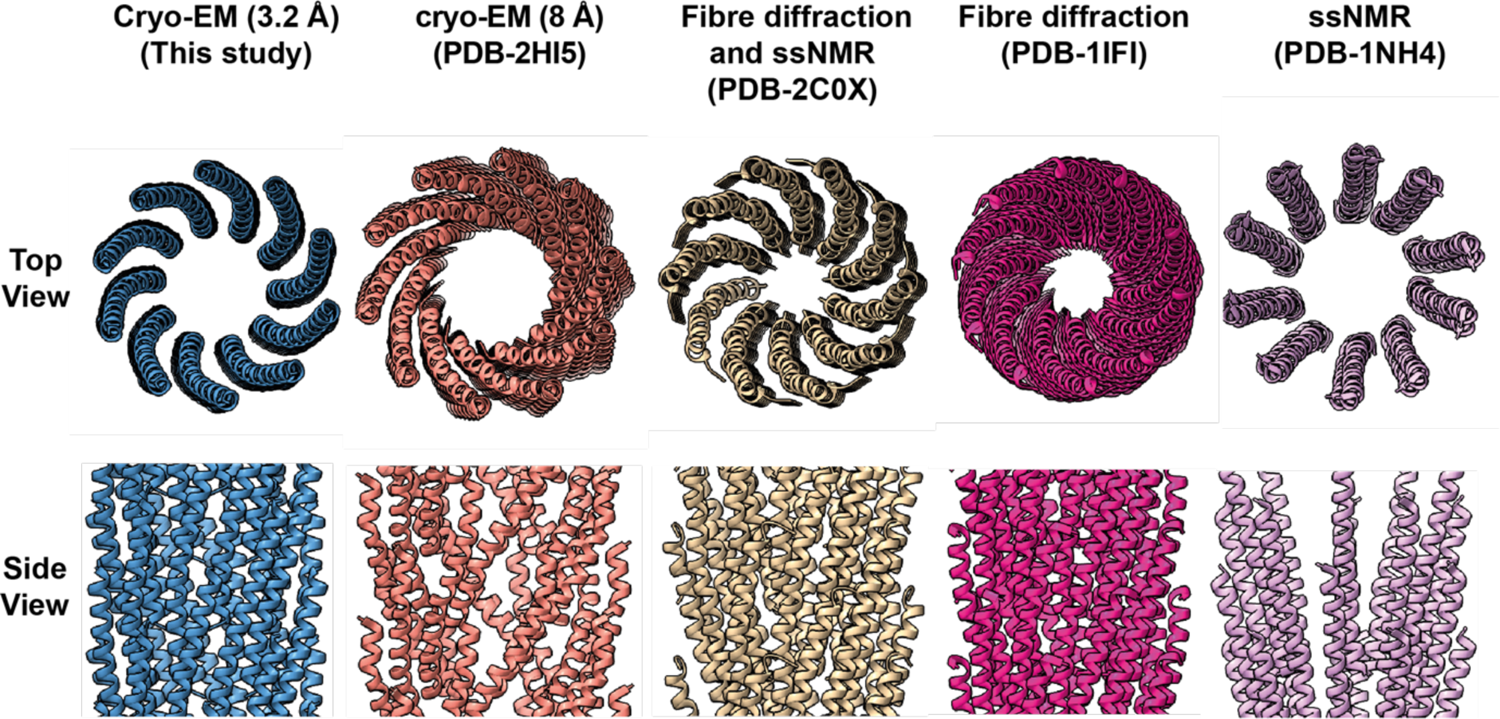
Comparison of the 3.2 Å-resolution cryo-EM structure obtained in this study with previous structural models of fd. Shown are ribbon diagrams (top and side view) of the 3.2 Å-resolution cryo-EM structure reported in this study compared to previously proposed fd capsid models based on an 8 Å-resolution cryo-EM map (PDB 2HI5), ssNMR and fibre diffraction combined (PDB 2C0X), fibre diffraction alone (PDB 1IFI), and ssNMR alone (PDB 1NH4).

**Figure S4:**
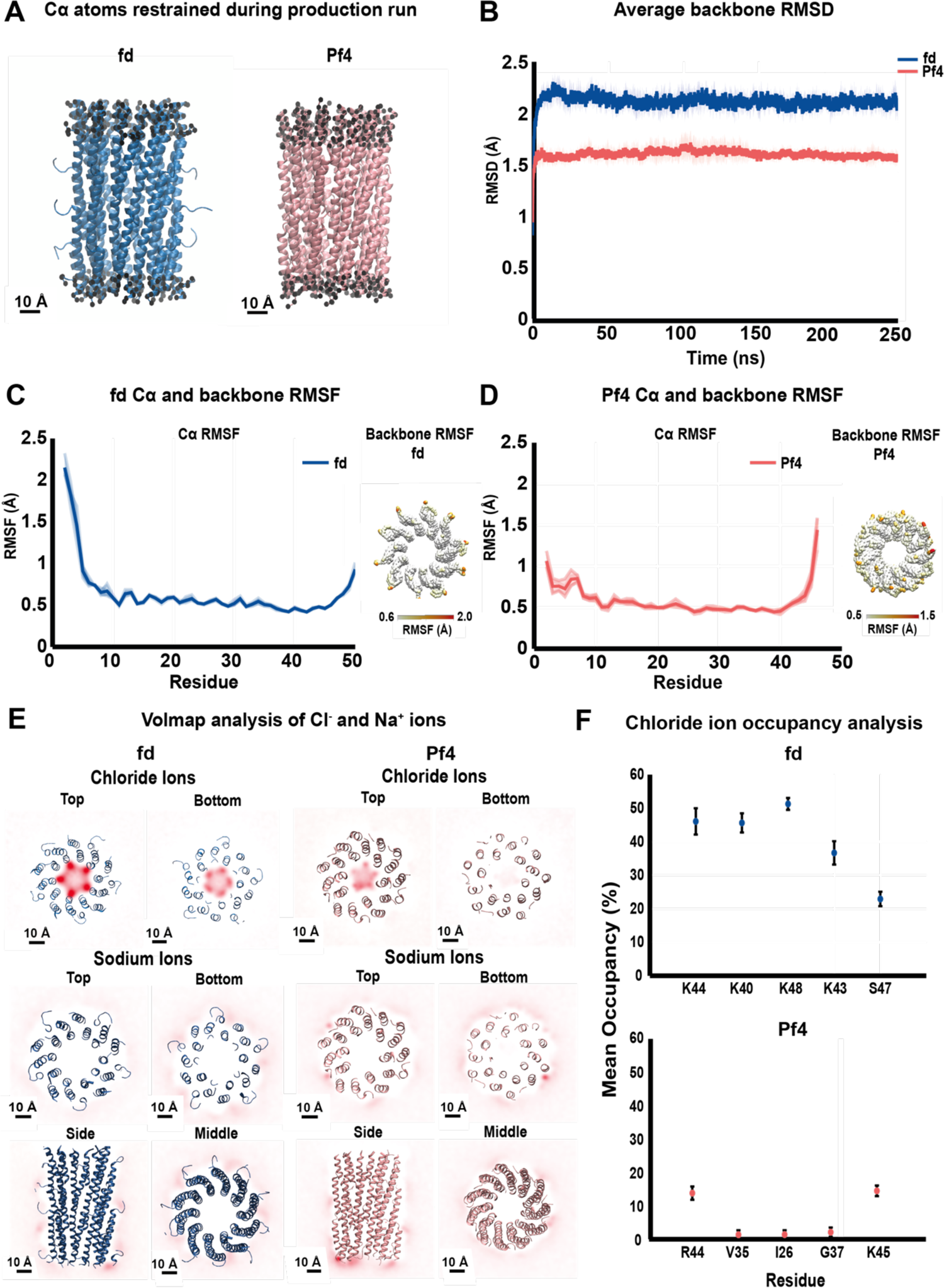
Atomistic molecular dynamics simulations of fd and Pf4 capsid proteins. **A)** Side view of fd (left) and Pf4 (right) showing C*α* atoms restrained during simulation production runs. Restrained C*α* atoms are shown as black spheres. **B)** Backbone RMSD plot of simulated systems for 250 ns in 0.15 M NaCl. Average RMSD (Å) is plotted, for fd (blue) and Pf4 (salmon) with the standard deviation plotted in transparent colour. Both simulation systems are stable. **C)** Fd C*α* RMSF plot (left) and backbone RMSF values plotted on the simulated structure (right) over 250 ns of simulation in 0.15 M NaCl. Average C*α* RMSF (Å) values are plotted in solid blue (left), with individual repeats plotted in transparent blue. RMSF show higher values at the N-terminus. **D)** Corresponding plots for Pf4, with average C*α* RMSF values plotted in salmon (left), with individual repeats plotted in transparent colour. RMSF values plotted on the simulated structure (right). **E)** Weighted ionic density of sodium and chloride ions (calculated using volmap analysis in the VMD software) averaged over all trajectory frames of fd (left) and Pf4 (right). Density of ions are shown in red. Dark red indicates higher ionic density. Volmap slice offsets used were 0.5 for side and middle, 0.28 for bottom and 0.67 for top views. There is a lack of sodium ions in the capsid lumen although they are present around the capsid on the outside. **F)** Quantification of chloride ion residue occupancy for positively charged C-terminal residues of fd (top) and Pf4 (bottom). Quantification was performed using PyLipID for ion analysis. Mean occupancy data is plotted, with error bars depicting SEM. The highest chloride ion occupancy residues are plotted for both systems, with overall higher occupancy detected for fd.

**Figure S5:**
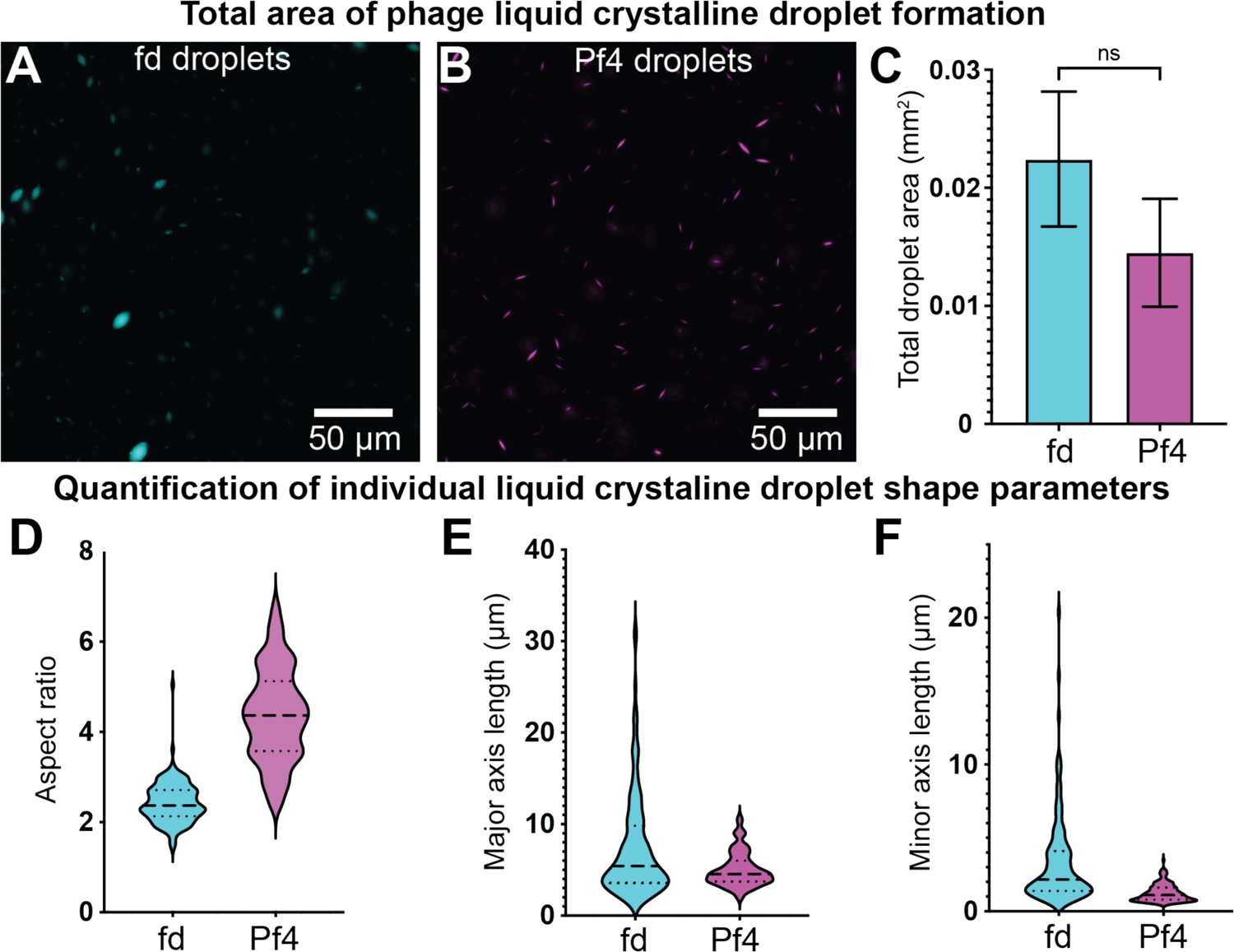
Quantification of phage liquid crystalline droplet morphology. **A-B)** Representative light microscopy images with comparable total liquid crystalline droplet area formed by A488-labelled fd (cyan) and Pf4 phages (magenta). **C)** Bar chart showing total droplet area as assessed by light microscopy followed by segmentation of droplets. Values shown are the mean of three independent experiments and error bars represent standard deviation. **D)** Violin plot of aspect ratios of individual droplets. **E)** Violin plot of major axis lengths of individual droplets. **F)** Violin plot of minor axis lengths of individual droplets.

**Figure S6:**
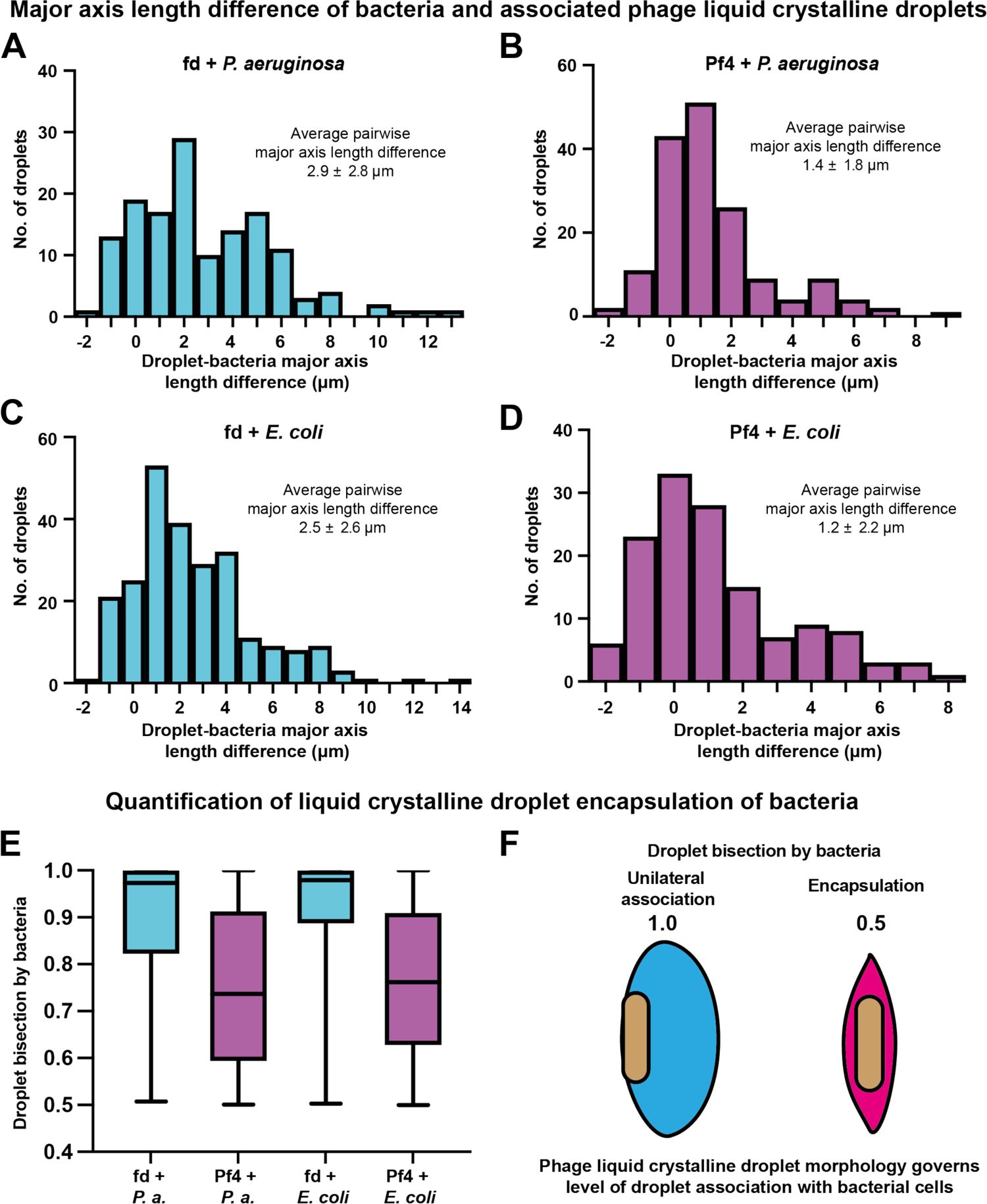
Quantification of phage liquid crystalline droplet association with bacterial cells. **A-D)** Histogram of the major axis length difference between bacterial cells and associated liquid crystalline droplet from semi-automated segmentation of images (Figure 5). Mean and standard deviation are reported from three independent experiments, **A)** fd + *P. aeruginosa* (n=144), **B)** Pf4 + *P. aeruginosa* (n=163), **C)** fd + *E. coli* (n=244) and **D)** Pf4 + *E. coli* (n=137). Fd + *P. aeruginosa* versus Pf4 + *P. aeruginosa*, **** P_value_ < 0.0001. fd + *E. coli* versus Pf4 + *E. coli,* **** P < 0.0001. Pf4 + *P. aeruginosa* versus Pf4 + *E. coli*, no significant difference. Fd + *P. aeruginosa* versus fd + *E. coli*, no significant difference. Pf4 + *P. aeruginosa* versus fd + *E. coli*, **** P_value_ < 0.0001. Fd + *P. aeruginosa* versus Pf4 + *E. coli*, **** P_value_ < 0.0001. **E)** Plot showing type of association of phage liquid crystalline droplets with bacterial cells from semi-automatic segmentation of images. This measurement was obtained by measuring the bifurcation of the major axis of the segmented bacterial shape of the liquid crystalline droplet. The ratio of the resulting two shapes was then calculated. **F)** Values of 0.5 indicate encapsulation of bacteria by droplets whereas values of 1 indicate unilateral association of droplets with bacteria.

## Supplementary Movie Legends

**Movie S1:** Cryo-EM map (10 *σ* away from the mean) and atomic model (ribbon depiction) of the bacteriophage fd.

**Movie S2:** Electron cryotomograms of liquid crystalline droplets formed by Pf4 (upper) and fd (lower).

## Supplementary information

**Table S1:**
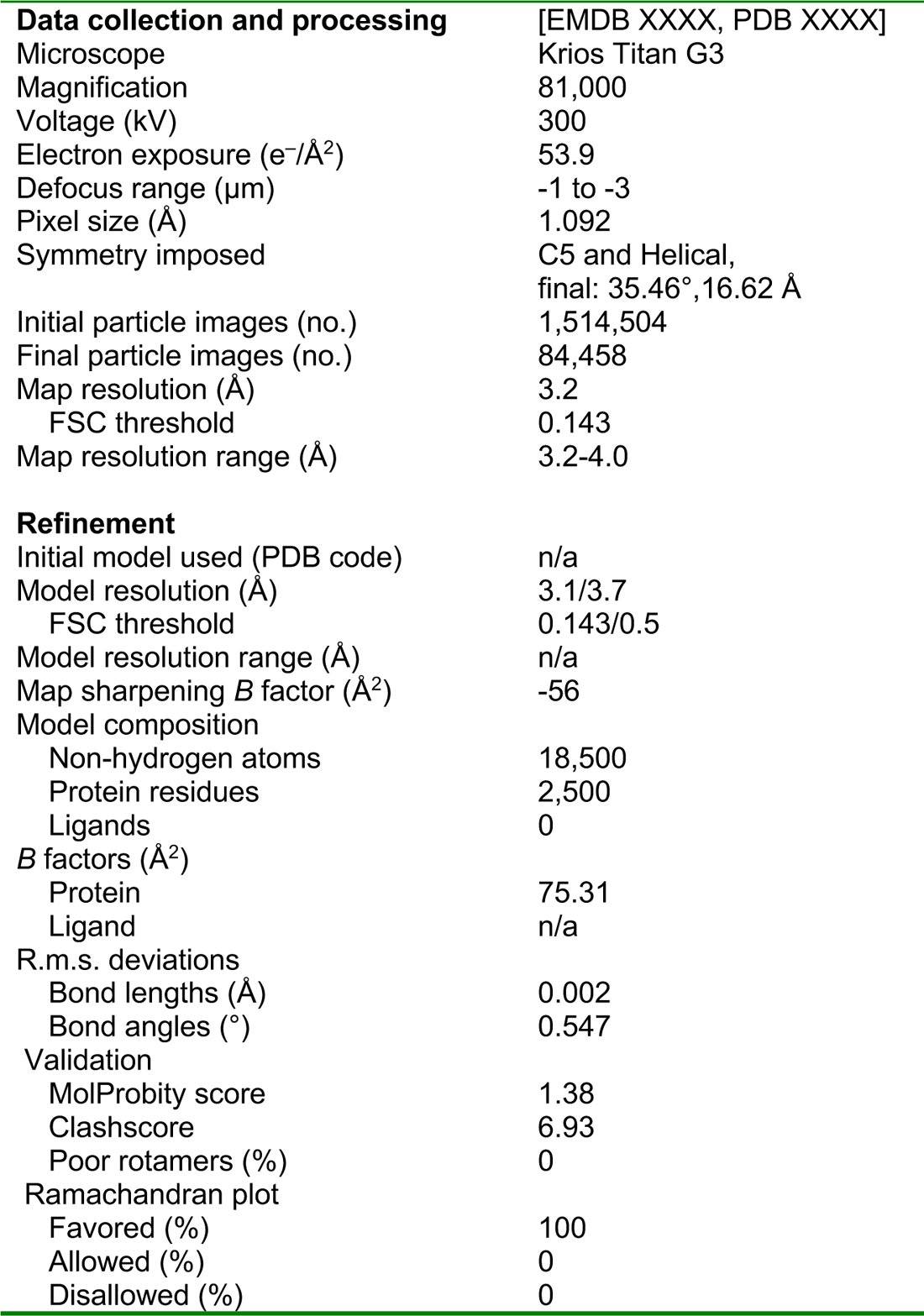
Data collection and processing statistics for fd capsid structure.

## Notes

### Competing Interest Statement

The authors have declared no competing interest.

